# The human LIN28B nucleosome is inherently pre-positioned for efficient binding of multiple OCT4s without H3 K27 acetylation

**DOI:** 10.1101/2023.10.06.559923

**Authors:** Tengfei Lian, Ruifang Guan, Bing-Rui Zhou, Yawen Bai

## Abstract

Pioneer transcription factors possess the unique ability to access DNA within tightly packed chromatin structures, playing pivotal roles in cell differentiation and reprogramming. However, their precise mechanism for recognizing nucleosomes has remained mystery. Recent structural and biochemical investigations into the binding interactions between the human pioneer factor OCT4 and the LIN28B nucleosome by Sinha et al.^1^ and Guan et al.^2^ have yielded conflicting results regarding nucleosome positioning, nucleosomal DNA unwrapping, binding cooperativity, and the role of N-terminal tail of OCT4. In this study, we undertook a comparative analysis of these two research efforts and delved into the factors contributing to the observed discrepancies. Our investigation unveiled that the utilization of human and Xenopus laevis core histones, along with a discrete two-step salt dialysis method, led to distinct positioning of DNA within reconstituted LIN28B nucleosomes. Additionally, our reanalysis of the electrophoretic mobility shift assay data showed that H3 K27 acetylation did not increase OCT4 binding to the internal sites of the nucleosome when normalized to input; instead, it promoted sample aggregation. Thus, the available experimental data support the notion that the human LIN28B nucleosome is pre-positioned for efficient binding with multiple OCT4s, and there is no compelling evidence for its regulation by histone modifications.

## Introduction

In a recent study, Sinha et al. ^1^ showed that OCT4 initially targeted the *LIN28B* DNA-containing nucleosome by interacting with the linker DNA and the N-terminal tail of histone H4, which led to the repositioning of DNA in the nucleosome that locates at multiple positions in its free form. OCT4 binding at the linker DNA was shown to be required for the efficient binding of two additional OCT4 molecules to the internal sites of the fully wrapped nucleosome using partial motifs. Sinha et al. also observed that H3 K27 acetylation influenced nucleosome positioning and enhanced OCT4 binding at the two internal sites, leading to the conclusion that histone modifications regulate OCT4 binding cooperativity.

In contrast, Guan et al.^2^ determined the structures of the *LIN28B* nucleosome and its complex with three OCT4 molecules at its internal sites. The DNA dyad position in Guan et al.’s nucleosomes differed by 13 base pairs (bp) from that observed in the nucleosome bound to a single OCT4 at the linker DNA, as reported by Sinha et al. Guan et al.’s data show that the free *LIN28B* nucleosome was already pre-positioned for efficient OCT4 binding at these three internal sites without the need of OCT4 binding at the liner DNA. One of these sites encompasses the full motif, leading to unwrapping of nucleosomal DNA. Moreover, Guan et al. demonstrated that OCT4s could work cooperatively to open H1-condensed nucleosome array containing the LIN28B nucleosome in the absence of H3 K27ac.

## Results and Discussion

What could be responsible for the discrepancies in the two studies? To investigate this question, we undertook a comparative analysis and found several differences in the sample preparation. For the nucleosome reconstitution, Sinha et al. used (i) *Xenopus larvis* core histones, (ii) 182 base pair (bp) DNA, and (iii) a discrete two-step salt dialysis method at 4 °C without subsequent heating and purification. As a result, the reconstituted nucleosome displayed two distinct bands in the native gel (see Figure 3b in Sinha et al.). Conversely, Guan et al. reconstituted the nucleosome using (i) human core histones, (ii) 162 (the original sequence used by Soufi et al.3) or 187 bp DNA, and (iii) a continuous salt-gradient dialysis with subsequent heating at 37 °C, for 3-5 hours, and purification. Guan et al. observed only single band for the nucleosome in the native gel. It’s worth noting that prior research had demonstrated that nucleosomes reconstituted with 5S DNA exhibited two bands without sample heating but only one band after heating.4 Consequently, the utilization of the two-step salt dialysis method at 4°C without heating may account for the nucleosome occupying different positions on the DNA.

For the reconstitution of the OCT4-nucleosome complex, Sinha et al. mixed the full-length OCT4, core histones and DNA together at 2M NaCl and then conducted two-step salt dialysis at 4 °C omitting any heating process. This approach, which simultaneously mixes OCT4, DNA, and core histones to reconstitute the OCT4-nucleosome complex for structural analysis, may potentially lead to a bias in favor of OCT4 binding to the linker DNA. Such a bias could kinetically trap the nucleosome structure at a local energy minimum (Figure 1). Furthermore, this method does not faithfully replicate the natural process of OCT4 targeting pre-formed nucleosomes in vivo.

**Figure 1.**
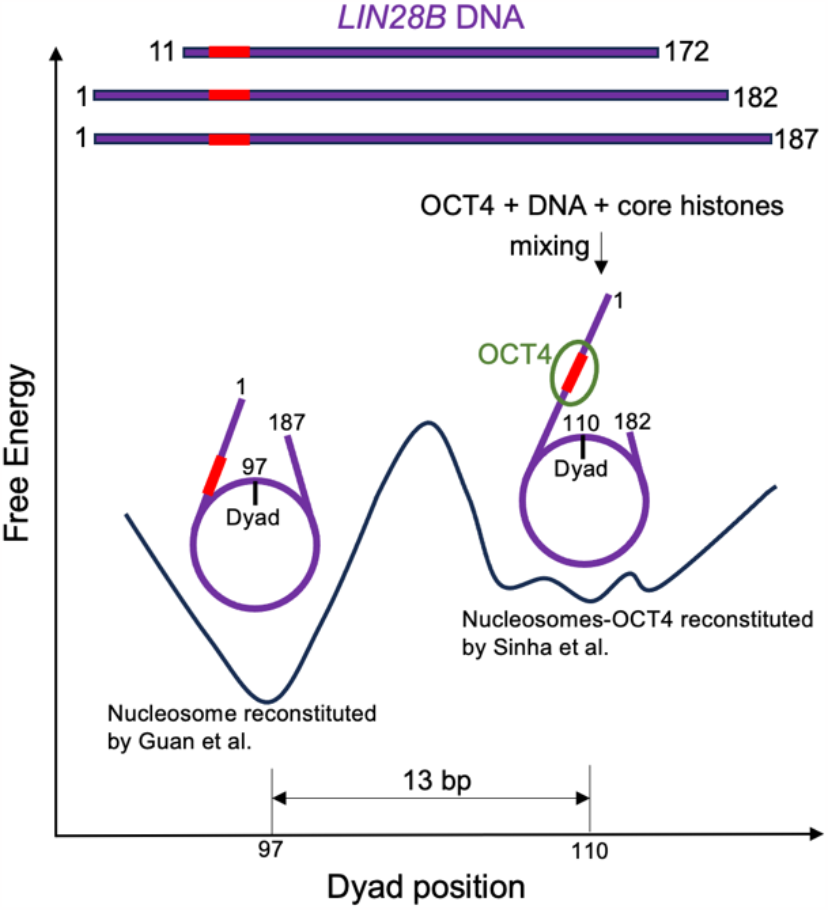
A proposed hypothetical energy landscape for the *LIN28B* nucleosome. The *LIN28B* DNA fragments used in the two studies include the initial 162 bp *LIN28B* DNA, the 182 bp DNA employed by Sinha et al., and the 187 bp DNA utilized by Guan et al. The OCT4 binding motif within the linker DNA region is denoted in red. The hypothesized energy landscape of the *LIN28B* nucleosome displays two distinct regions that are separated by a significant kinetic barrier. The nucleosome reconstituted by Guan et al. adopts a lower energy state and is well-positioned with the dyad at nucleotide 97 (PDB IDs: 7U0J and 8DK5 for the DNA fragment with162 and 187 bp, respectively). The nucleosome-OCT4 complex reconstituted by Sinha et al. is situated at a local energy minimum (PDB ID: 8G8G), which is positioned a few base pairs away from others in their reconstituted nucleosomes.

In contrast, Guan et al. introduced the maltose binding protein (MBP)-fused OCT4 DNA binding domain region (MBP-^DBDR^OCT4) to the pre-reconstituted nucleosome. Notably, Guan et al. demonstrated that MBP-^DBDR^OCT4 and MBP-^full-length^OCT4 exhibited the same binding affinity to the LIN28B nucleosome, while MBP alone did not interact with the nucleosome. Furthermore, the nucleosome structures observed in Guan et al.’s study consistently maintained the same DNA positioning, whether OCT4 was present or not. It is worth highlighting that Guan et al. employed a single-chain antibody fragment (scFv) to aid in their structural determination. Importantly, scFv does not interact with the nucleosomal DNA. Therefore, it is improbable that scFv binding could induce a 13 bp shift in the nucleosomal DNA. Additionally, the unwrapping of nucleosomal DNA, as evidenced in the structure of the OCT4-nucleosome-scFv complex, was independently confirmed through MNase digestion and FRET experiments, even in the absence of scFv.

The 187 bp DNA employed in Guan et al.’s structural study encompasses the 182 bp DNA region used in Sinha et al.’s research, albeit with a mutation in the linker DNA region^2^. This observation implies that the length of the DNA should not inherently influence the positioning of the nucleosome core particle. To further investigate the impact of DNA length in both studies, we conducted a nucleosome reconstitution experiment utilizing Sinha et al.’s 182 bp DNA, along with human core histones and their two-step salt dialysis method. The results showed two distinct bands on the native gel (Figure 2**a**). Upon denaturing gel analysis, it became apparent that the upper band contained approximately 50% less H2A compared to the lower band. This finding strongly suggests that the upper band represents hexasomes, while the lower band corresponds to octasomes (nucleosomes). It’s worth noting that in our hands, the human core histone octamer displayed lower stability. It is expected to observe a significant presence of hexasomes in our reconstituted sample. However, what came as a surprise was the observation of only one band for the octasome (nucleosome) in contrast to the two bands seen in Sinha et al.’s study. A closer examination of the nucleosome structure revealed differences between human and Xenopus laevis core histones at 24 specific positions, some of which are near the DNA (see Figure 2**b** and Supplementary Fig. 1). These findings strongly suggest that the choice between human and Xenopus laevis core histones could result in distinct positioning of the LIN28B nucleosome.

**Figure 2.**
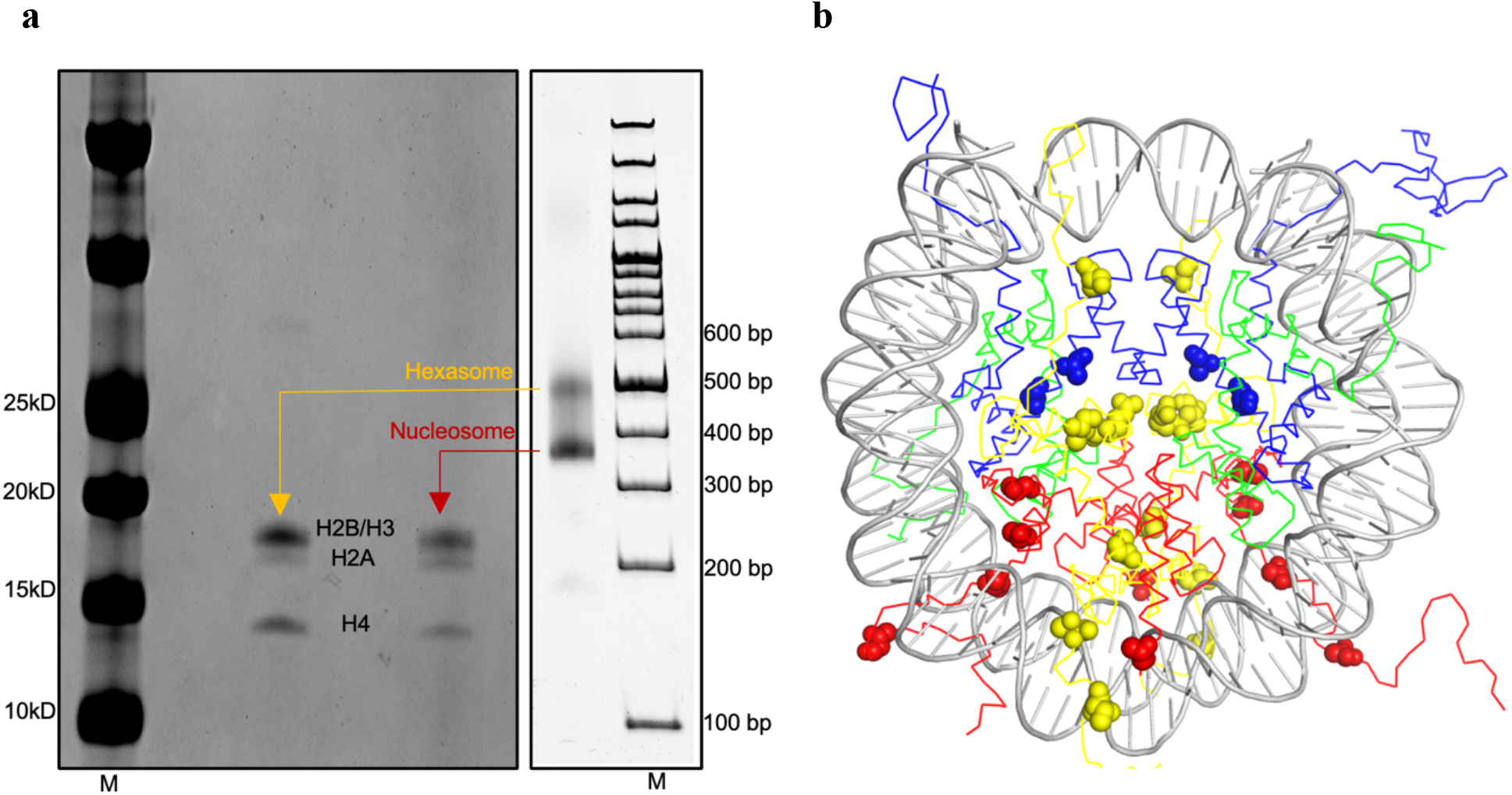
Nucleosome reconstitutted using human core histones and the two-step salt dialysis method and illustration of residues different in human and *Xenopus larvis* core histones. **a**. SDS gel (left) of the two bands from the native PAGE gel (4%) of the reconstituted nucleosome using the two-step dialysis method and 182 bp as in Sinha et al. The gel experiments were repeated three times. The ratios of the intensity of H2A over the intensity of all histone bands ([I^H2A^]/[I^total histones^]) are 10.0 ± 0.9% for the upper band and 19.1 ± 0.8 % for the lower band, respectively. **b**. Illustration of the *Xenopus Larvis* core histone residues (spheres) (H2A in yellow, H2B in red, H3 in blue, and H4 in green) that are different from those in human core histones at the corresponding positions in the structure of the nucleosome core particle (pdb ID: 1kx5).

Sinha et al. concluded that the N-terminal tail of OCT4 played a role in impeding nucleosome packing and condensation of nucleosome arrays. However, their biochemical assays employed Mg^2+^ to induce mono-nucleosome packing and condensation of nucleosome arrays, instead of using linker histone H1, which is known to regulate linker DNA length and nucleosome packing in human chromatin.^5,6^ Additionally, their analysis is based on the widely-held perception that the H4 tail interacts with the acidic patch of neighboring nucleosomes in condensed nucleosome arrays, but there is no direct evidence for this claim.^7^ In contrast, Guan at al. have shown that deletion of both the N- and C-terminal tails of OCT4 does not affect its binding affinity for the nucleosome. Moreover, our biochemical experiments demonstrated that the DNA binding domain region of OCT4 alone, without the N- and C-terminal tails, can unwrap the nucleosome, displace H1, and locally open the H1-condensed nucleosome array containing the *LIN28B* nucleosome.

Sinha et al. also proposed that H3 K27 acetylation releases its interactions with the linker DNA, facilitating nucleosome repositioning and the binding of OCT4 at the internal sites. However, their conclusion of nucleosome repositioning based on changes of MNase-seq pattern (Sinha et al.’s Figure 3**e, g** and Extended Data Fig. 9**j**) is questionable. For example, it is possible that H3 K27ac simply alters the local DNA environment near the entry region, facilitating MNase cleavage at nearby locations. In their model, they assume that OCT4 interacts with DNA motifs at OBS2 and OBS3 sites within a fully wrapped nucleosome. However, they provide no structural or biochemical evidence demonstrating that the nucleosome remains fully wrapped when OCT4 binds to internal sites. Furthermore, it’s important to note that full-length OCT4 has a high propensity for aggregation. Their electrophoretic mobility shift assay revealed significant nucleosome and OCT4 aggregation at higher OCT4 concentrations, with H3 K27 acetylation exacerbating this aggregation (Figure 3**e** in Sinha et al. and Supplementary Fig. 2). Consequently, the apparent increase in OCT4 binding at internal sites relative to linker DNA, attributed to H3 K27 acetylation, may potentially be influenced by the same factors leading to aggregation, such as OCT4-OCT4 interactions. Indeed, we find that H3 K27 acetylation does not lead to an increase in OCT4 binding at internal sites when normalized to the initial nucleosome input (Supplementary Fig. 1). It appears that H3 K27 acetylation primarily promotes nucleosome aggregation in the presence of OCT4, possibly due to increased hydrophobicity. Notably, the C-terminal tail of OCT4 is known to contribute to the formation of condensates in vivo by interacting with Med1. ^8^ Therefore, using full-length OCT4 protein to study OCT4-LIN28B DNA nucleosome interactions without Med1 may not be the ideal choice, given its propensity for aggregation. Our experiments with a more soluble form of full-length OCT4 fused to MBP demonstrate efficient binding to internal sites even in the absence of H3 K27ac (Supplementary Fig. S1**A** in Guan et al.). Lastly, a recent study has shown that H3 K27 acetylation is not essential for chromatin accessibility and enhancer activity in mouse embryonic stem cells.^9^

In brief, the discrepancies in nucleosome positioning observed between the research conducted by Guan et al. and Sinha et al. can be primarily attributed to the use of core histones from humans versus Xenopus laevis, as well as variations in the methods employed for nucleosome reconstitution. There is no substantial evidence to support the notion that H3 K27 acetylation is a prerequisite for enhancing the cooperative binding of human OCT4 by altering nucleosome positioning. The LIN28B nucleosome, comprising human core histones, is inherently prepositioned for the efficient binding of multiple full-length OCT4 proteins when fused to MBP to enhance solubility.

## Acknowledgments

We thank Dr. Ken Zaret for helpful comments and Dr. Mario Halic for discussion. This work is supported by the intramural research program at the Center for Cancer Research, National Cancer Institute, National Institutes of Health.

## Author contributions

T.L. conducted the experiments. Y.B. wrote the commentary with input from T.L., R.G., and B-R.Z.

## Declaration of interests

Authors declare no competing interests.

## Supplemental Methods

### Expression and purification of human core histones

Recombinant human core histones H2A, H2B, H3, and H4 were expressed individually in E. coli BL21(DE3) cells as described in a previous study. Briefly, E. coli cells harboring each histone expression plasmid were grown at 37 °C in 2 x YTB Broth. 0.3 mM IPTG was added to induce recombinant protein expression for 3 h at 37 C, When OD600 reached around 0.6–0.8. The cells were harvested and resuspended in 50 mL of buffer A (50 mM Tris-HCl, 500 mM NaCl, 1 mM PMSF, 5% glycerol, pH 8.0), followed by sonication on ice for 60 min. The cell lysates were centrifuged at 20,000 x *g* for 20 min at 4 °C. The pellet containing histones was resuspended in 50 mL of buffer A and 7M guanidine hydrochloride. The samples were rotated for 12 h, and the supernatant was recovered by centrifugation at 96,000 x *g* for 60 min at 4 °C. The supernatants were dialyzed against buffer C (5 mM Tris-HCl, pH 7.4, 2 mM 2-mercaptoethanol, 7 M urea) for three times. The supernatant was loaded to Hitrap S column chromatography (GE Healthcare). The column was washed with buffer D (20 mM sodium acetate, pH 5.2, 100 mM NaCl, 5 mM 2-mercaptoethanol, 1 mM EDTA, and 6 M urea). The histone protein was eluted with a linear gradient of 100–800 mM NaCl in buffer D. The purified histones were dialyzed against water for three times, and freeze-dried.

### Preparation of 182 bp LIN28B DNA

The 182 bp LIN28B DNA fragment with the same sequence as that used in Sinha et al.’s study was prepared by PCR amplification using the 187 bp LIN28B DNA fragment in Guan et al.’ study as the template, followed by ethanol precipitation and purification using the POROS column. The PCR products were pelleted by 75% ethanol containing 0.3 M NaAc at pH 5.2. The sample was incubated for 60 min at -20 °C, followed by centrifugation. The pellet was resuspended in TE buffer. The sample was loaded to POROS column chromatography (GE Healthcare). The column was washed with buffer containing 20 mM Tris-HCl, pH 7.4, 5 mM 2-mercaptoethanol, and the DNA was eluted by a linear gradient of 0–2 M NaCl.

### Reconstitution of the 182 bp LIN28B nucleosome with human core histones

Purified recombinant human core histones in equal stoichiometric ratio were dissolved in 6 mL unfolding buffer (7 M guanidine-HCl, 20 mM Tris-Cl at pH 7.4, 10 mM DTT) and subsequently dialyzed against refolding buffer (10 mM Tris-Cl at pH 7.4, 1 mM EDTA, 5 mM b-mercaptoethanol, 2 M NaCl) for about 12 h. The mixture was centrifuged at 20,000 x *g* to remove any insoluble material. Soluble octamers were purified by size fractionation on a Superdex 200 gel filtration column. Nucleosome assembly was done by following the ‘double bag’ dialysis method of Sinha et al. Briefly, the histone octamer and the 182 bp DNA fragment were mixed in equimolar ratios in high salt buffer containing 25 mM HEPES (pH 7.5), 2 M NaCl and 2 mM DTT. The mixture was filled into a 3.5kDa cut-off dialysis bag and dialyzed against the high salt buffer for 3 hours. The dialysis bag was then transferred into 1L of the buffer containing 25 mM HEPES (pH 7.5), 1 M NaCl and 1 mM DTT for overnight dialysis. The buffer was changed to 25 mM HEPES (pH 7.5) and 1 mM DTT for the second overnight dialysis. The reconstituted nucleosome was stored at 4 °C for the EMSA study.

### Electrophoretic assays and band intensity measurement

The reconstituted nucleosome samples were analyzed using native polyacrylamide gel electrophoresis (4%) in 0.2 x TBE at 120 V for 70 min at 4 °C. After electrophoresis, the gel was stained with ethidium bromide (EtBr). To investigate the histone components of the bands in the native gel, the bands were cut out and soaked in 2x SDS loading buffer. After heating at 90 °C for 10 mins, the samples were loaded to the SDS page gel. Band intensities in the gels were quantified using ImageJ.

**Supplementary Fig. 1.**
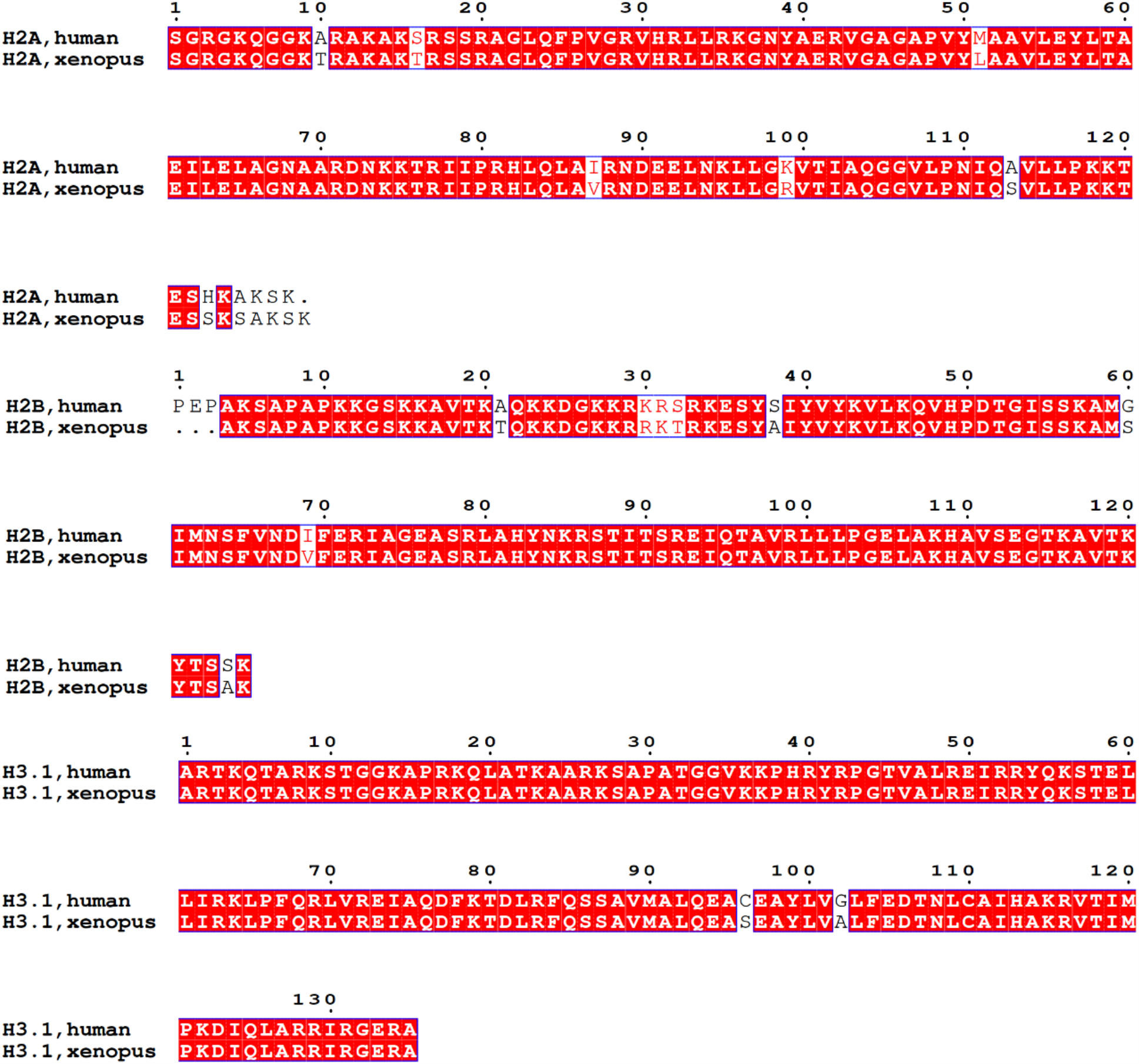
Comparison of human and *Xenopus larvis* core histone sequences.

**Supplementary Fig. 2.**
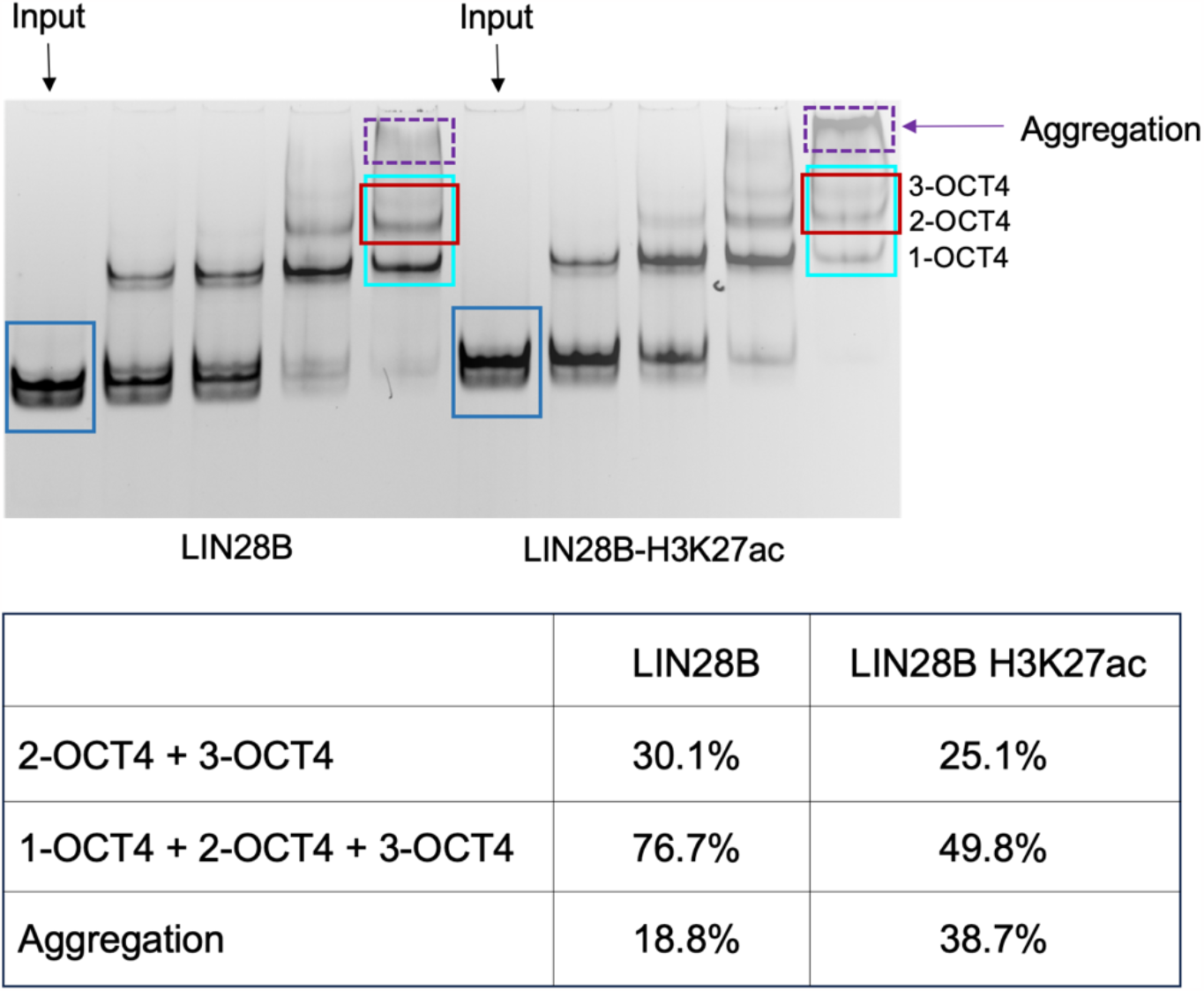
H3 K27 acetylation does not increase the population of the nucleosome bound to two and three OCT4s versus input. Illustration of band intensity measurement of OCT4 binding to the LIN28B nucleosome using Sinha et al.’s gel (Fig. 3**b** in Sinha et al.^1^) and ImageJ (upper panel; the boxes indicate the area used for intensity measurement), and percentage of the intensity ratio of the nucleosome bound to OCT4 over the input.

## Notes

### Competing Interest Statement

The authors have declared no competing interest.

